# Exploratory analysis and error modeling of a sequencing technology

**DOI:** 10.1101/043042

**Authors:** Michael Inouye, Kerrin S. Small, Yik Y. Teo, Heng Li, Nava Whiteford, Tom Skelly, Irina Abnizova, Daniel J. Turner, Panos Deloukas, Dominic P. Kwiatkowski, Clive G. Brown, Taane G. Clark

## Abstract

Next generation DNA sequencing methods have created an unprecedented leap in sequence data generation, thus novel computational tools and statistical models are required to optimize and assess the resulting data. In this report, we explore underlying causes of error for the Illumina Genome Analyzer (IGA) sequencing technology and attempt to quantify their effects using a human bacterial artificial chromosome sequenced to 60,000 fold coverage. Seven potential error predictors are considered: *Phred* score, read entropy, tile coordinates, local tile density, base position within read, nucleotide call, and lane. With these parameters, logistic regression and log-linear models are constructed and used to show that each of the potential predictors contributes to error (P<1×10^−4^). With this additional information, we apply the logistic model and achieve a 3% improvement in both the sensitivity and specificity to detect IGA errors. Further, we demonstrate that these modeling approaches can be used as a feedback loop to inform laboratory methods and identify specific machine or run bias.

## Introduction

With the increasing pace of DNA sequencing technology it is crucial to understand the limitations and error biases of these technologies to both ensure the quality of what is produced and to expand what is reasonably achievable (Olson, 2002). Central to this tenet is the identification of the causes of error and an assessment of their quantitative effect. With these parameters characterized, statistical models can be built to maximize the accuracy of the resulting DNA sequences and eliminate error’s underlying cause.

Many ‘next-generation’ sequencing technologies have been or will be available shortly (Bentley, 2006,Braslavsky, et al, 2003,Margulies, et al, 2005,Shendure, et al, 2005). Currently, three of the most popular are available from Roche Bioscience, Applied Biosystems, and Illumina, Inc. and will be utilized in the upcoming 1000 Genomes Project. For the Roche (454) Genome Sequencer, which uses the pyrosequencing method (Ronaghi, et al, 1998), Brockman et al. have sought to calculate accurate *Phred* quality scores (Ewing, et al, 1998) in order to create a common language for all sequencing-by-synthesis and Sanger sequencing methods (Brockman, et al, 2008). They rightly note that this is of crucial importance to platform analyses and statistical modeling, and that base calling errors cause problems in both read alignment and single-nucleotide polymorphism (SNP) calling.

Herein, we empirically assess the causes of error for the Illumina Genome Analyzer (IGA) sequencing platform, using a bacterial artificial chromosome (BAC) of human DNA which has been both capillary sequenced and run across eight IGA machines to a total depth of 60,000 fold coverage. Exploratory analysis techniques are used to identify and characterize the main causes of error. Using these parameters, we construct and validate a logistic regression model to optimize error prediction. With the model, we are able to substantially improve the quality of IGA reads by accurately identifying specific base errors in each read. Furthermore, the model can be tailored to maximize performance on specific IGA machines and potentially, given a suitable control sample, individual runs. By using its *β* coefficients (odds ratios), we also demonstrate that the model can act as a guide to minimize machine variation and as a feedback loop for laboratory methodologies.

## Results & Discussion

### IGA imaging, base calling, and filtering

Sample loading for IGA sequencing occurs on a flowcell which has eight parallel lanes, each containing amplified fragments of DNA on approximately 300 imaging areas, or “tiles”. From lane one through eight, the flowcell is iteratively scanned as it is put through N cycles of sequencing chemistry, each extending the complementing DNA polymer by one base.

By default, IGA provides an informatics pipeline to handle the data coming from the tile imaging process. This pipeline is made up of three modules: Firecrest, Bustard, and Gerald. See **Figure 1** for an overview. Briefly, imaging of the tile for each cycle results in four images, one for each specific nucleotide fluorophore. Firecrest then performs a background correction and identifies clusters of amplified molecules on each tile, after which occurs a crosstalk correction which compensates for the overlapping frequency response of the labeled nucleotides (Li, et al, 1999). A phasing estimation is applied that corrects for those molecules which do not incorporate a nucleotide successfully or happen to incorporate more than one. Bustard implements the phase correction and calculates base calls for each cluster as a function of the corrected signal. Gerald identifies and filters overlapping clusters using these signals, and finally outputs the base calls and IGA quality scores in the form of fastq files (see **Methods**). Implementation of SEAM is done after Gerald.

**Figure 1:**
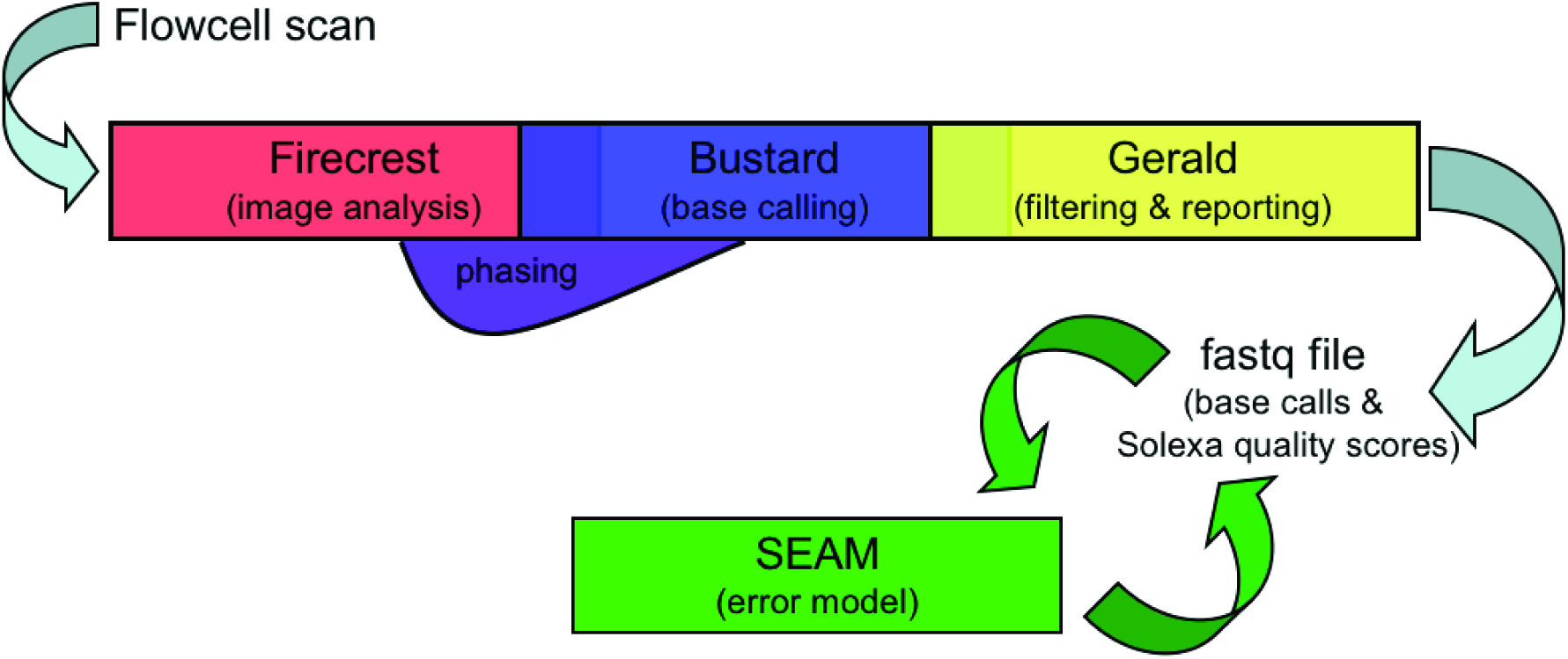
IGA analysis pipeline overview. After tile images are collected from scanning the flowcell, they enter three analysis modules: Firecrest, Bustard, and Gerald. The Firecrest module performs image analysis to identify cluster intensities, correct crosstalk for overlapping dye florescence, and estimate phasing corrections to compensate for unsuccessful nucleotide incorporation or unblocked read molecules. Bustard implements the phase correction and performs base calling using maximum post-correction intensity, while Gerald filters overlapping read clusters and reports the base calls and IGA quality scores. Currently, SEAM is implemented after Gerald using the resulting fastq files.

### Exploratory analysis of error predictors

In order to model the IGA sequencing platform, we consider seven potential predictors of error:

1. Read sequence entropy (*V*_*w*_)
2. *Phred* score
3. Base position within the read
4. Nucleotide calling bias
5. Tile coordinates
6. Tile cluster density
7. Lane

Typically, the error control of sequencing data, e.g. weighting by *Phred* score, is entangled with the method of alignment, of which many algorithms exist (Li H., et al. 2008,Hillier, et al, 2008,Li R., et al, 2008,Ning, et al, 2001,Smith, et al, 2008). However, it is advantageous to have more complex modeling techniques exist outside of alignment algorithms so that (a) more alignment algorithms can use them and (b) we can be confident that the errors we see are attributable to the technology and not a particular alignment tool.

To achieve both of these aims, it is necessary to address the somewhat circular nature of training the error model since there is currently no known way to identify errors other than by alignment. For (a) above, the training and validation of the error model using one alignment algorithm (as many sequencing centers use) will result in a model which corrects for technology biases but whose coefficients (in this case, the *β* coefficients) also incorporate alignment artifacts. This will be advantageous if one uses an algorithm exclusively and is only interested in optimizing data quality with the given technology. For (b), we require a more general model which can control for reads which cause alignment artifacts. We can then create a feedback loop in which the modeling coefficients of a machine or a run accurately inform laboratory methodologies and instrument design in addition to acting as a normalizing factor for data already generated. Thus, our approach is to define a metric which can identify regions likely to give discordant alignments for short-read algorithms. We utilize a well-known method, Shannon entropy (Shannon, et al, 1949,Valdar, 2002), and modify it with a sliding window to give us a measurement of the innate sequence complexity of a read (see **Methods**). We then stratify our analyses to separate areas of homopolymers and other repetitiveness, which have been well known to bias alignment results. We also consider entropy as a parameter in the error model.

The *Phred* score as calculated from IGA quality scores (see **Methods**) is an imperfect measurement of accuracy. **Figure 2a** shows a plot of observed versus predicted *Phred* score for each of eight machines with the best possible fit along the *x* = *y* axis (average coefficient of determination, *R*^*2*^, of 0.961). With increasing deviation of an observed from an expected *Phred* score, one can expect a loss of sensitivity or specificity. In addition, variability in the performance between machines can cause differential bias if data from >1 machines is merged. In our experiment, we observe noticeable variability across eight machines, with sample variance of *R*^*2*^ = 0.000128. When taking into account an observed frequency distribution of each *Phred* score, a weighted calculation of *R*^*2*^ gives an average of 0.965 with greater machine variation (sample variance = 0.000958). **Figure 2a** also demonstrates IGA’s tendency to underestimate error probability at both low (<11) and high (>33) *Phred* scores and in some cases actually showing decreasing accuracy as quality score increases. In the context of SNP calling, this may result in an inflation of false positives. However, it should be noted *Phred* scores and IGA quality scores are asymptotically related and this may not be fully captured by the transformation between the two, thus explaining part of the deviation we see. The most elegant solution would be re-evaluating the signal to noise ratios, but error models should be able to correct for this as well.

**Figure 2:**
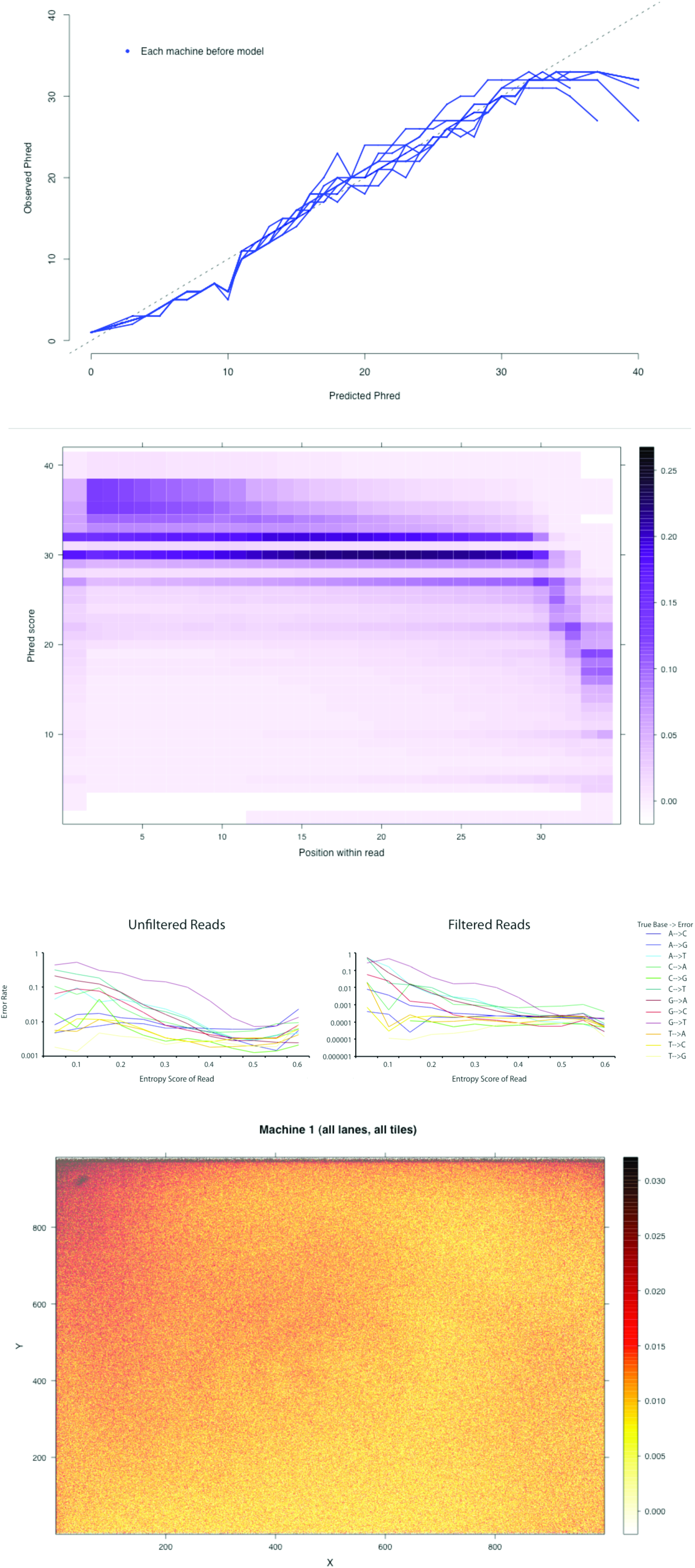
Predictors of IGA sequencing errors. In **(a)**, for each *Phred* score the observed probability of error was calculated. This probability was then converted back into an observed *Phred* score which was then plotted against the predicted. Perfect correlation is the dotted line. We also observe variability across machines. The decline of base quality within a read is shown in **(b)**. At each base position in a read we calculate the frequency distribution of observed *Phred* score, which we use as a surrogate for the signal to noise ratio. This shows the probability of a base calling error increasing as position increases with an apparent inflection point at approximately 30 bases. Errors are also not uniformly distributed across nucleotides (c). In particular, T for G and A for C substitutions are inflated for both unfiltered and filtered data (*Phred* > 20 and Maq alignment score > 40). Part **(d)** gives an aggregate view of error rate across a tile. For each X,Y coordinate on all tiles from machine #1, we calculate the proportion of errors out of the total base calls. These were then mapped out on a virtual tile and each proportion colored (the sidebar). An analogy would be stacking all 300 tiles on top of one other and “seeing” the average error by looking down the long axis. From our observations, each machine had a different pattern of error across the tile, however the tile edges were consistently more error-prone.

With many sequencing-by-synthesis methods, a base call’s signal to noise ratio varies depending on its position within a read, thus causing each read to have a non-uniform quality distribution. As a proxy for the signal to noise ratio distribution, we assessed *Phred* score distribution as a function of base position within a read (**Figure 2b**). Generally, *Phred* scores decrease as base position within a read increases following a somewhat bimodal distribution with scores of 30 and 32 arising with disproportionate (20-25%) frequency. Curiously, the first base appears to be of similar quality as a base from about the 15^th^ position, a possible artifact of reagent dead volumes, component temperatures, and/or initial laser power.

Since imaging of each tile is largely dependent on the fluorescence of different chemically labeled nucleotides, we can hypothesize that each nucleotide is called with a different efficiency and, since scanning is an iterative process, the error bias of each nucleotide is dependent on the preceding nucleotide. To minimize the alignment bias which homopolymeric and repetitive sequences can introduce, we calculate the probability of observing different nucleotide substitutions as a function of read entropy (**Figure 2c**). We observe almost an order of magnitude difference between nucleotide substitution rates for the most prevalent entropy range 0.45 – 0.55, highlighting the importance of incorporating nucleotide bias into an error model. In addition, there exists a consistently elevated substitution rate (0.6% overall) of A for C (A/C) and an entropic dependence for the T/G substitution (1.2% overall) (**Supplementary Fig. E**). The latter may partially be explained by the presence of Gs in or directly adjacent to many homopolymeric T regions in the BAC. To assess the effect of preceding nucleotides further, we plot the above for dinucleotides, the second of which is substituted (**Supplementary Fig. G**). We find that an A/C substitution is independent of the preceding base while a TT/TG substitution is the most prevalent. The A/C substitution is also overrepresented with regard to *Phred* score, read position, and lane in both unfiltered and filtered (only bases with *Phred* >20 and reads >40 Maq alignment score are considered) data. However, the TT/TG substitution becomes more frequent as lane number increases (lane 8 having the most substitutions), potentially due to a relatively greater susceptibility to free radical build up during imaging. In our experience, this can be corrected for in the laboratory by using an imaging buffer with greater ascorbate protection (data not shown), implying that as light exposure time increases less light-mediated radicalization will occur. It should be noted that the wavelengths which fluoresce G and T are emitted from the same laser and have overlapping emission/excitation bands (similarly for A and C), thus one will expect there will always be some level of T/G and A/C bias.

With GA1 optics, imaging of the tile also exhibits a spatial bias with different areas of the tile experiencing different probabilities of error (**Figure 2d**). In general, the closer a read is to the edge of the tile, the greater the likelihood that any base call within the read will be an error. Additionally, spatial patterns of error are also lane dependent with the last lane, lane eight, experiencing noticeably worse performance (**Supplementary Fig. A**). In conjunction with our observation that T/G substitutions occur most frequently, **Supplementary Fig. A** also suggests that the increasing lane error, especially in lane eight, is almost wholly attributable to these substitutions. Related to this spatial tile dependency is the interference of neighboring reads with a focal read (also known as cluster overlap). As the distance between reads on a tile decreases, the more difficult it becomes to resolve each base call. IGA’s Gerald module implements a cluster overlap correction or “purity” filter, however we further investigate resolution by calculating the local read density of every unfiltered read on each individual tile (**Supplementary Fig. B**). At low and medium density, there appears to be no effect on the error rate distribution; at high density, we did observe an increasing shift in the error distribution, however this is very unlikely to have a large effect since densities of >0.05 reads/coordinates^2^ occur with a frequency of 3.2×10^−7^.

Using these seven predictors as parameters, we adopt logistic regression and log-linear frameworks to assess the relationship between predictors in the same model and the additional predictive ability of models which utilize all potential predictors as compared to models incorporating just *Phred* or alignment scores.

### Model design and performance

All potential predictors considered individually in the logistic and log-linear models showed evidence of being associated with errors (P<1×10^−4^). When all predictors are included in logistic and log-linear models, their effects are homogeneous across machines (**Supplementary Tables B & C**). For the logistic model, the strongest predictors were *Phred* score and entropy (*V*_*w*_), with single unit increases leading to reduced risks of error. In particular, univariate and multivariate models both showed that an increase in *Phred* score of one reduced the risk of error by ~25% (**Table 1**); this effect also appears to be approximately linear (**Figure 3a**), although deviations of the predicted *Phred* from the data cause this relationship to become more nonlinear. Similarly, the effect of *V*_*w*_ on the (log) error risk also appears to be linear (**Figure 3b**), with an increase of 0.01 units leading to a reduced risk of ~1%. Logistic models which incorporate the other potential predictors and *Phred* score had consistently greater area (~1.5%) under their Receiver Operating Characteristic (ROC) curves and a ~1.1% improvement in calibration, thus reflecting a greater predictive ability overall (see **Figure 3c** for ROC curves of machine 8, which is typical of other runs). In addition, after transforming the new predicted probability of an error into *Phred* form, the average coefficient of determination with the observed *Phred* score was 0.988 across all machines. Simply using the given *Phred* score and filtering those calls with scores <20 to predict error does have reasonable sensitivity and specificity (both ~0.90), however there are additional gains in predictive ability from considering *Phred* score as a continuum and using the other predictors with both sensitivity and specificity (both ~0.93), see **Table 1**. Validation of the model on data from machines that were not used to train the model led to slightly diminished predictive ability. However, these differences are small due to the homogeneity of coefficients across machines.

**Figure 3:**
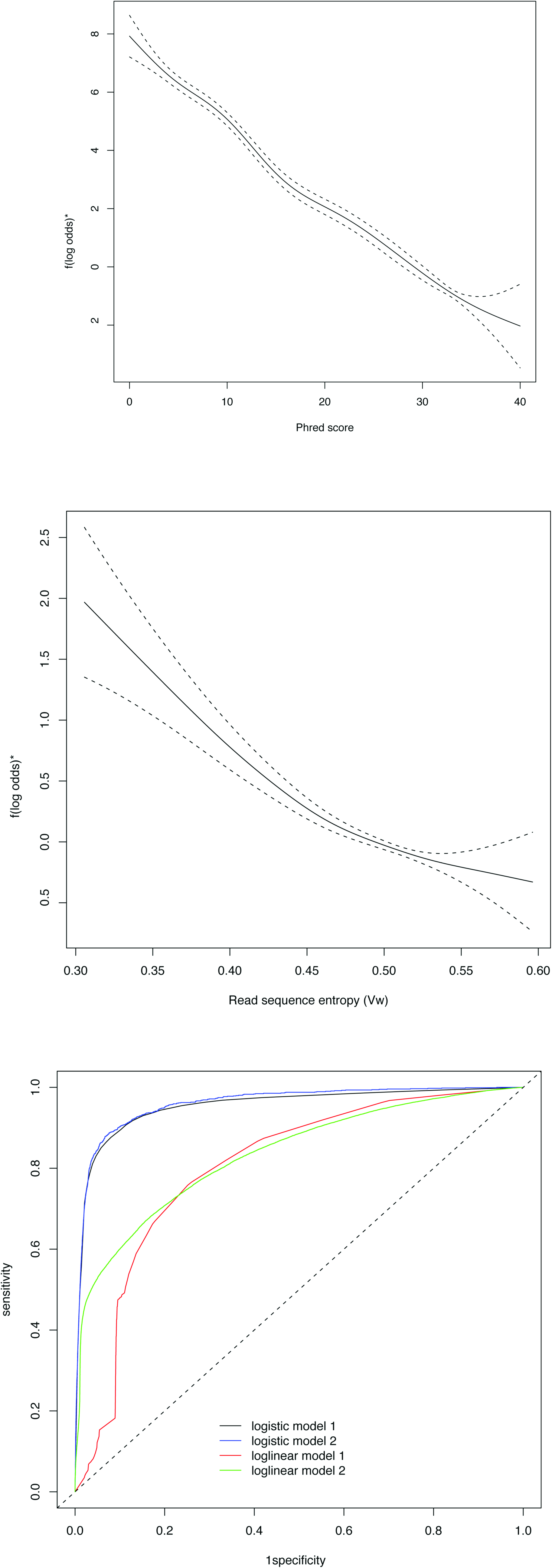
Application of SEAM. For (a) and (b) we present the relationship between the log-odds of an error and *Phred* and read entropy, *V*_*w*_, respectively. *These were estimated by fitting splines within a generalized additive model. In (c), we show ROC curves for four models: a logistic model with *Phred* only (logistic model 1), a logistic model with all predictors (logistic model 2), a log-linear model with sequence alignment score only (log-linear model 1), and a log-linear model with all predictors (log-linear model 2).

**Table 1:** Logistic model performance

| Machine | Model 1 |  |  |  |  | Model 2 |  |  |  |  |  |  |
| --- | --- | --- | --- | --- | --- | --- | --- | --- | --- | --- | --- | --- |
|  | Risk estimates* | Specificity** | Sensitivity** | AURC | RMSE | Risk estimates* | Specificity** | Sensitivity** | AURC | RMSE | Val AURC*** | Val RMSE*** |
| 1 | 0.741 | 0.903 | 0.883 | 0.932 | 0.093 | 0.737 | 0.931 | 0.923 | 0.948 | 0.092 | 0.943 | 0.102 |
| 2 | 0.763 | 0.860 | 0.892 | 0.935 | 0.084 | 0.764 | 0.911 | 0.930 | 0.952 | 0.083 | 0.947 | 0.094 |
| 6 | 0.753 | 0.860 | 0.908 | 0.947 | 0.093 | 0.752 | 0.909 | 0.935 | 0.952 | 0.092 | 0.949 | 0.099 |
| 8 | 0.762 | 0.886 | 0.908 | 0.945 | 0.095 | 0.760 | 0.912 | 0.934 | 0.955 | 0.094 | 0.953 | 0.104 |

Model 1 consists of *Phred* score only, and Model 2 consists of position, call, $V_w$ , lane, density, radial distance, and *Phred*. AURC = area under the ROC curve; P = p-value; Risk estimates are odds ratios; RMSE = root mean square error; \* all p-values < 1e-100; \*\* median of best combination of sensitivity and specificity, \*\*\* median based on predictions from models trained on other machines.

Implementation of the log-linear model had indicated that *V*_*w*_, radial distance, density, lane, and alignment score (AS) were all strong predictors of the number of errors (**Supplementary Table C**). However, the predictive ability of the model was inferior to the logistic regression approach. The relationship between Maq’s AS and logarithm of the expected number of errors is not linear (**Supplementary Figure H**), but overall an increase of one unit of AS leads to a 3% reduction in the risk of an error. It should be noted however that this particular modeling approach may, in fact, be more useful to implement in conjunction with a particular alignment algorithm since it can incorporate any form of alignment score.

In using the modeling coefficients (odds ratios) as more practical guides, we observe the difference in nucleotide calling bias between machines, a potential indicator of suboptimal optics calibration. As can be seen from **Supplementary Table B**, machines 1, 6, and 8 exhibit significantly more bias than machine 2. In conjunction with a noticeably greater lane effect, it can be hypothesized that the bias in machine 1 may be compounded by either increased laser exposure or problematic imaging buffer. These observations provide educated first guesses for the causes of machine variability and can lead the trained sequencer to correct or compensate in the laboratory. In addition, with the pooled use of a suitable control sequence, the coefficients can be used at the intra-machine level to monitor and normalize individual runs by training SEAM on the control sample and transferring the resulting coefficients to the test sample.

### Implications and Improvements of SEAM

SEAM’s logistic regression model adds accuracy and reproducibility to IGA base calls and runs, which translates to improvements in downstream applications. Besides SNP calling, short read assembly is particularly sensitive to sequencing errors; one example being the use of De Bruijn graphs which currently attempt to identify errors via topological features of the graph, like the formation of two largely redundant paths or “bubbles” (Zerbino, et al, 2008). Unfortunately, this approach cannot as yet make use of the features of a particular run or lane which should only reduce the frequency of bubbles. One particularly attractive application is quantitative expression profiling via cDNA sequencing (Nagalakshmi, et al, 2008,Wilhelm, et al, 2008). Since alignment bias will affect quantitative expression patterns, it is essential to minimize machine errors and misalignment, however these measurements are still subject to IGA’s filtering of overlapping read clusters. With the percentage of reads filtered at approximately 40%, this introduces a large amount of uncertainty if not bias, which could potentially arise if overlapping (or non-overlapping) read clusters are sequence dependent. In the absence of supporting quantitative measures, it is therefore crucial that future filters are able to dissect overlapping clusters and laboratory protocols are modified to minimize their frequency.

SEAM’s use of IGA’s *Phred* calculation is largely a surrogate for signal to noise ratio, which is currently handled by the Bustard module. A far more elegant input parameter would be the signal to noise ratio itself; it follows then that SEAM could be integrated into the IGA informatics pipeline around Bustard (instead of after the Gerald outputs), thus, with better error detection, some overlapping clusters could be differentiated (although assumedly with an elevated proportion of errors which still makes alignment difficult). Leaps in the technology and instrumentation may also limit the effectiveness of an error model. These leaps will inevitably introduce new parameters which should be taken into account while, ideally, minimizing previous parameter(s) propensity for error. However, the more precise error estimates with gains in sensitivity and specificity provided by a successful error model can also minimize the divide between old and new technology, making data more amenable to combined analyses. Additionally, implementation of technology gains is not uniform for all laboratories or sequencing centers, thus it is important to understand the differential bias arising from them. Finally, it is possible to implement more complex statistical methodologies than presented here. An improvement to the model itself would be to apply a Bayesian logistic regression (Clark, et al, 2007), but this would necessarily be more computationally intensive. Alternatively, it is possible to implement a machine learning approach, such as that using support vector machines that update parameter estimation and predictions with increasing data (Cristianini, et al, 2000). Further research is being conducted in this area.

## Methods

### IGA sequencing and read alignment

DNA for sequencing came from the human BAC, bCX98J21 (Shizuya, et al, 1992). DNA was prepared by Illumina and sequenced at the Wellcome Trust Sanger Institute using the Solexa sequencing technology platform (GenomeAnalyzer I or II, Illumina) following standard protocols. Briefly, 5μg of bCX98J21 BAC DNA was sheared randomly by nebulisation with compressed nitrogen at 35psi for 6 minutes. Fragmented DNA was purified using a QiaQuick spin column and end-repaired using T4 DNA polymerase and Klenow polymerase with T4 polynucleotide kinase to generate blunt ended fragments with phosphorylated 5’ termini. Following a second column purification, a 3’ A-overhang was created using a 3’–5’ exonuclease-deficient Klenow fragment. DNA was again purified with a QiaQuick column, and Illumina paired-end adapter oligonucleotides were ligated on. The DNA was size-selected by performing agarose gel electrophoresis and stabbing a scalpel blade into the gel at 290bp, so as to obtain 200bp inserts. DNA was washed from the blade with 10mM Tris-HCl pH8.5 and enriched for fragments with Solexa primers on both ends by an 18-cycle PCR reaction.

Following quantification of amplified DNA on an Agilent Bioanalyser 2100, paired-end flowcells were prepared for each machine on the supplied cluster station according to the manufacturer’s protocol, using DNA at a final concentration of 4pM. Clusters of PCR colonies were then sequenced on each GenomeAnalyzer using supplied protocols. Images from the instrument were processed using the manufacturer’s software.

Each run was analyzed using the Illumina GA Pipeline version 0.3x, default settings were used, this includes “purity” filtering which discards those reads with potentially ambiguous calls. The purity of a read is calculated as the minimal *p* value over the first 12 bases where *p* is defined as:

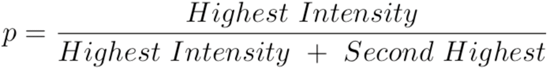

Any read which is not pure (*P* > 0.60) is discarded. Approximately 40% of reads are removed by this filter; cluster summary statistics for the runs used here are shown in **Supplementary Table A**.

Read alignment to the BAC capillary sequence was performance using the Maq algorithm (Li H., et al. 2008) and software version 0.6.3, freely available at http://maq.sourceforge.net under a GNU public license. Alignment of each run was done using Maq default parameters and follows a standard protocol (cite http://maq.sourceforge.net/maq-man.shtml). Read map positions, quality scores, etc. were extracted via Maq’s internal functions, mapview and pileup. For each machine, there was an average of 7,914x coverage at each reference base.

### Refinement of the BAC capillary sequence

With relatively deep coverage, one can make accurate judgments about the base call in a reference sequence. This was done via a simple frequentist approach at each base in an alignment using all runs. For each strand, the probability of observing the reference base call was calculated and compared to that of the other three base calls. If on both +/-strands the probability of observing the reference base is less than any of the other bases, then the reference base was deemed a suspect call. This approach clearly identified four bases (reference call/IGA call) of >99% mismatch within 30 bp of each other at BAC positions 169520 (C/T), 169532 (C/A), 169542 (G/A), and 169547 (A/G) as errors in the reference sequence. All further analyses of these four bases consider the IGA call to be the true base. Three other base positions (5451, 5453, and 114642) were deemed suspect calls, however upon inspection these were obvious alignment errors due to their position within homopolymeric runs and elevated read coverage.

### Sequence entropy and inference of machine error

To assess sequence complexity, entropy is a common measure (Valdar, 2002). We introduce a modified Shannon entropy (*V*_*w*_) which utilizes a sliding window of variable size to capture as many sub-states as possible of the system, a read sequence. Traditionally, Shannon entropy is defined as

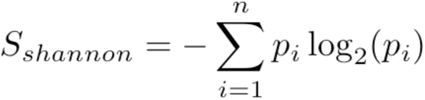

where *p*_*i*_ is the observed frequency of *i*, and *n* is the number of possible states of *i*. Here, the normalized entropy of a read is defined as

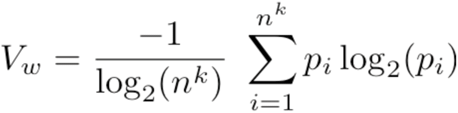

where *n* is 4 (possible nucleotide calls), *k* is the number of bases in window *i*, and *p*_*i*_ is the observed frequency of *i*. The normalizing term is the maximum possible entropy at a given *k*, or log_2_(4^*k*^). Since we are examining very short reads of <40 bp, we unfortunately have too few measurements to accurately determine entropies at large *k*, thus we restrict our calculations to *k* = 1, 2. The minimum *V*_*w*_ for a read across all *k* is taken to be its sequence complexity. This captures areas of repetitiveness across the BAC (**Supplemental Fig. C & D**).

### Phred and Pseudo-Phred scores

For each base call, the Bustard module uses the signal-to-noise ratio to calculate a IGA quality score which is asymptotically related to *Phred* (Ewing, et al, 1998), thus it can be thought of as a pseudo-Phred score. If the error probability of a base call is *p*_*e*_ and *ψ* is the IGA quality score, then

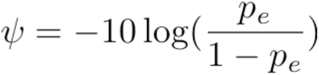

Transformation of *ψ* into *Phred*, *ϕ*, and its relation to *p*_*e*_ are as follows:

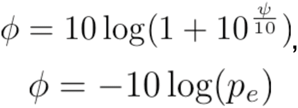

It is *ϕ* on which we perform our analyses. To assess the accuracy of *ϕ* in predicting the actual error rate, we calculated the probability of observing an error at a given *ϕ*, converted it to the corresponding *Phred* score, then plotted these observed *Phred* scores versus the predicted values. These are stratified to show inter-machine variation (**Figure 1a**).

### Effect of intra-read base position

As with other sequencing-by-synthesis methods, IGA experiences a drop in quality as base position within a read increases. This is almost exclusively due to a decreasing signal to noise ratio as imaging of the read progresses, however alignment algorithms can also have an effect (e.g. by assigning heavier weights to earlier bases). In order for an error model to be useful in read alignment, it must concentrate on signal to noise ratio therefore we investigated the effect of read position on *Phred* score (**Figure 2b**). For each position, we calculated the frequency of each observed *Phred* score and plotted these score distributions as a function of their position.

### Nucleotide bias

To determine the extent to which errors are biased toward nucleotides, we calculated the substitution rates for each nucleotide and, being particularly sensitive to alignment, further stratified this by each read’s sequence complexity. In observing bias in the T for G and A for C substitutions, it was hypothesized that this might be dependent on IGA run lane, read position, and *Phred* score (**Supplementary Fig. F**).

The possibility of photo-interference or differential fluorescence decay led us to calculate the above for dinucleotides as well (**Supplemental Fig. G**).

### Effect of tile coordinates and cluster density

The probability of error was calculated as a function of each read’s tile coordinates and was then stratified by machine and lane (**Supplementary Fig. A**). The potential clustering of reads with similar sequence complexities (e.g. homopolymeric regions) was also assessed as a possible explanation for tile edge effects, none was observed.

Given the potential for imaging biases, the probability of error as a function of local read density on the tile was investigated. For each read on each tile, the local read density was calculated as

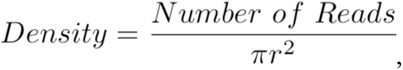

the number of reads within a defined coordinate radius of the focal read divided by the area. To get a suitable range of measurements within the smallest radius, we chose the coordinate radius as ten (πr^2^ = 314.159 coordinates^2^). The probability of observing an error given a local read density was then determined; since each density has a probability of observing a certain error rate, a heat map was constructed to show these probability distributions (**Supplementary Fig. B**). To avoid elevated error rates on tile edges, we excluded all reads within 100 coordinates of a tile edge.

### Statistical approaches for error modeling

We adopted two statistical modeling strategies: (a) a logistic regression approach to predict the odds of an error in a read at a particular base and given a set of other characteristics of the read, and (b) a log-linear modeling approach to estimate the number of errors in a read given a set of read characteristics. For method (a), the model is

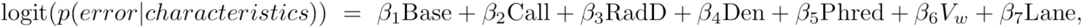
where logit(p) = p/1-p, *Base* refers to its location within the read, *Call* is the IGA call, *RadD* is the read’s radial distance from the center of the tile, *Den* is the read’s local density on the tile, *Phred* is the quality score, *V*_*w*_ is sequence complexity, and *Lane* is the read’s lane on the flowcell. The coefficients (log odds) were estimated using iteratively re-weighted least squares (IRWLS) (McCullagh, et al, 1989), but it is also possible to estimate these with a Bayesian framework (Clark, et al, 2007). Note that we have simplified the notation above, as categorical variables (e.g. *Lane*) are represented by a number of variables (e.g. seven lane variables comparing to a baseline group to represent the eight possible lanes). The predicted probability of an error, *p.error*, is

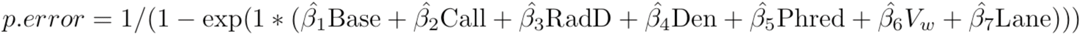

where the *β* hats are estimated as described above.

For method (b), the model is

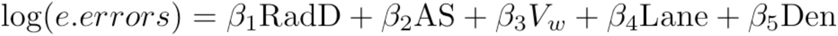

where AS is the alignment score and the other variables are defined above, and the coefficients (risk ratios) are also estimated using IRWLS. The expected number of errors, *e.errors*, is

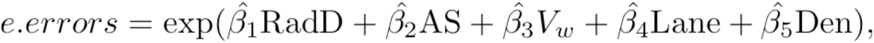

where the *β* hats are estimated as described above, and the expression can be used to threshold for one or more errors.

### Model Validation

We assessed the linearity of the relationship between the risk of errors and the continuous predictors considered (e.g. *Phred* score) by fitting splines within a generalized additive model (Harrell, et al, 1996). For each machine run we generated 10 datasets consisting of 3,000,000 randomly chosen reads. Validation was performed by training both models on single runs, and using the resulting coefficients to predict the results for other runs. Two measures of predictive ability were applied: the area under the ROC curve (AURC) and the root mean squared error (RMSE) (Hastie, et al, 1990). The ROC curve is a function of the sensitivity and specificity, where sensitivity is the proportion of true errors that are correctly identified and specificity is the proportion of true negatives that are correctly identified by our approach. An AURC equal to 0.5 indicates random predictions and a value of 1.0 corresponds to a perfectly discriminating model. In the case of the log-linear model, we estimated the AURC by dichotomizing the number of errors into whether there are zero or greater than zero errors. The RMSE indicates whether the predicted error probabilities agree with observed error probabilities, and in the case of the log-linear model we compared the expected and observed number of errors.

## Acknowledgements

We thank Richard Durbin for thoughtful discussions and Kathryn Holt for critical reading of the manuscript. Funding for this work came from the Wellcome Trust, the Bill and Melinda Gates Foundation, and the Medical Research Council UK.

